# Beyond suppression: a stand-off effect of mitigation measure reduces seabird bycatch in pelagic longline fisheries

**DOI:** 10.64898/2026.11.25.769318

**Authors:** Daisuke Ochi, Noriyosi Sato, Nobuhiro Katsumata, Hiroshi Minami

## Abstract

Bycatch in longline fisheries is one of the causes of population decline in albatrosses and other pelagic seabirds worldwide, and tori-lines and weighted branch lines are the mitigation measures most widely deployed to address this threat. Despite their widespread use, it remains unclear exactly how these devices reduce bycatch and whether combining them provides added protection beyond either device alone. Using experimental longline operations conducted in the western North Pacific with two albatross species commonly caught in the region, this study linked seabird attack behavior at the point of capture with the actual outcome of bycatch, revealing an overlooked mechanism. Deploying double tori-lines, rather than single, did more than reduce attacks near the vessel: it pushed birds away entirely, keeping them clear of baited hooks before capture could occur. Weighted branch lines, by contrast, reduced attacks but did not push birds away, and for one species even increased the likelihood that attacks occurred within reach of the vessel. This “stand-off effect” of a double tori-line, rather than attack suppression alone, was the strongest predictor of reduced bycatch, and its benefit did not simply add on top of the bycatch reduction by weighted branch lines. The two albatross species also responded differently to the same devices, reflecting differences in their foraging behavior. Conservation and fisheries managers designing bycatch mitigation programs should therefore evaluate candidate measures against the behavior of the seabird species actually exposed to local fishing grounds, rather than relying on standard specifications developed for different regions and species.

## 1. Introduction

Fisheries bycatch is a lethal event for vulnerable seabirds and it ultimately poses serious conservation risks for their populations. Large numbers of seabirds have been killed through such fishery interactions; reducing incidental capture is therefore a component of sustainable fisheries management at both regional and international levels (Phillips et al., 2016; Dias et al., 2019). The development of technology to mitigate seabird bycatch in pelagic longline fisheries remains a conservation priority. Fatal bycatch in longline fisheries occurs primarily during line setting, when seabirds attempting to take the bait become hooked or entangled, are pulled underwater as the baited line sinks, and drown. Consequently, technology has been developed to prevent seabirds from accessing the hooks (Brothers et al. 1999; Domingo et al. 2017; Robertson et al. 2018).

Tori-lines and weighted branch lines are very popular, and regarded as the most effective and fundamental mitigation measures for pelagic longline fisheries together with night setting (Løkkeborg 1998; Clarke et al. 2014; Domingo et al. 2017). Tori-lines, also known as streamer lines, consist of a mainline fitted with deterrent streamers, and are towed from vessel stern with a rigid pole during line setting. Tori-lines have gained acceptance among fishers because of their ease of installation and low cost relative to other mitigation devices. Mitigation effectiveness is also thought to increase when two tori-lines are deployed simultaneously, one on each side of the vessel (Sato et al. 2012; Melvin et al. 2013, 2014) , and when sufficient aerial coverage is maintained (Sato et al. 2013), and maintaining a sufficient aerial coverage by tori-line (Sato et al. 2016; Ochi et al. 2022). Weighted branchline increases sink rate, carrying baited hooks more quickly to depths beyond the reach of seabirds (Robertson et al. 2010; Clarke et al. 2014; Lee et al. 2026). Combining tori-lines with weighted branch lines allows hooks to sink below the reach of birds during the period in which tori-lines are effective (Melvin et al. 2014).

Despite this evidence supported by empirical verification, development and evaluation of varied tori-line designs have continued without consensus on how, and to what extent, they reduce seabird bycatch, and the absence of a stable performance indicator constrains the design of evidence-based specifications. The most direct indicator is a comparison of bycatch counts under standardized experimental conditions (Sato et al. 2012; Melvin et al. 2014; Jiménez et al. 2020), but this approach requires accepting that seabird mortality will occur in each experiment, raising conservation concerns when experiments are repeated. In addition, once a device reaches sufficient effectiveness, bycatch events become rare except in areas of high seabird density, which limits quantitative evaluation. An alternative approach is to count seabird attacks on baited hooks during line setting (Cortés and González-Solís 2018; Gilman et al. 2021), but this metric has been used as a proxy for bycatch risk without verification that attack frequency is linked to bycatch. Gaps remain in understanding how these devices—tori-lines in particular—function. The extent to which tori-lines reduce attacks near the vessel, and whether this reduction translates into fewer bycatch events, remains unclear. The effectiveness of weighted branch lines when used alone has not been evaluated.

A further gap concerns the extent to which behavioral responses to mitigation devices differ among seabird species. Previous studies have suggested that differences in foraging behavior among species within a regional seabird assemblage produce differences in device effectiveness and susceptibility to bycatch (Murray et al. 1993; Jiménez et al. 2020; Lucas and Berggren 2022). In mitigation device development, foraging behavior differences are often characterized by diving capability (Frankish et al. 2021; Bell et al. 2025), but behavior at the sea surface—including approach pattern, boldness toward bait, and competition over bait—also varies, not only between families and genera but within the same genus (Zhou and Brothers 2022). Such differences should manifest as differences in behavioral response to mitigation devices. Clarifying species-specific differences provides information for selecting mitigation measures suited to the seabird assemblage in a given fishing region, and the combined behavioral-observation and bycatch-based framework described above allows this to be tested. Studies on the effectiveness of tori-lines and weighted branch lines have mostly been conducted in the southern hemisphere under high seabird attack pressure (Melvin et al. 2014; Sato et al. 2016; Domingo et al. 2017). Seabird assemblages in the North Pacific differ from those in the south in species composition and behavior (Sato et al. 2012, 2013; Melvin et al. 2014; Gilman et al. 2021). Whether device combinations effective in southern fisheries perform similarly in northern waters remains unresolved, particularly for weighted branch lines.

This study used data from experimental longline operations conducted in the North Pacific to evaluate seabird bycatch mitigation measures, applying Bayesian modeling to assess attack behavior during line setting and resulting bycatch events jointly. Based on the model results, we examined how tori-lines, weighted branch lines, and their combinations contribute to bycatch mitigation, and discuss evaluation indices and mitigation strategies for seabird bycatch, and to assess how bycatch mitigation measures should be optimized for local seabird communities.

## 2. Materials and methods

### 1) Operational experiment

The experiment was carried out aboard a chartered longline vessel, *F/V Takei-Maru No. 2* (196 tons, 37 m in total length), from December 2011 to June 2012. A total of 74 longline operations were conducted in the western North Pacific off Japan (26–38°N, 143– 148°E). Twelve operations in which no seabirds were observed were excluded, leaving 62 operations for analysis.

The longline configuration followed the Japanese shallow-setting style targeting swordfish (*Xiphias gladius*) and blue shark (*Prionace glauca*). Lines were set in the afternoon before dusk and hauled starting at dawn. Each operation deployed 960 hooks (240 baskets of four branch lines between floats) at a vessel speed of 7 knots and a target fishing depth of 40– 70 m. Buoy lines were 8 m long. Baited hooks were cast by hand astern of the vessel on the port side beyond the propeller turbulence. The mainline was set at 3.3 m s⁻¹ by a line shooter, and the branch lines and buoys were deployed every 7 s. Bait consisted of whole, fully thawed chub mackerel (*Scomber japonicus*).

The tori-lines used in the experiment were of the same design as those used in previous studies in the southern hemisphere (Melvin et al. 2013) and North Pacific (Sato et al. 2012). Each line was 150 m long and consisted of three sections, long-streamer, short-streamer, and underwater towing section (Fig. 1, Table 1). Tori-lines were attached to 6.5 m fiberglass poles extending 3–4 m outboard from both sides of the vessel, approximately 10 m above the sea surface. When only a single tori-line was deployed, only the port-side pole was used. To prevent damage to the tori poles caused by entanglement with fishing buoys, a break point consisting of a small metal swivel was inserted at the junction between the short-streamer and underwater towing sections.

**Figure 1.**
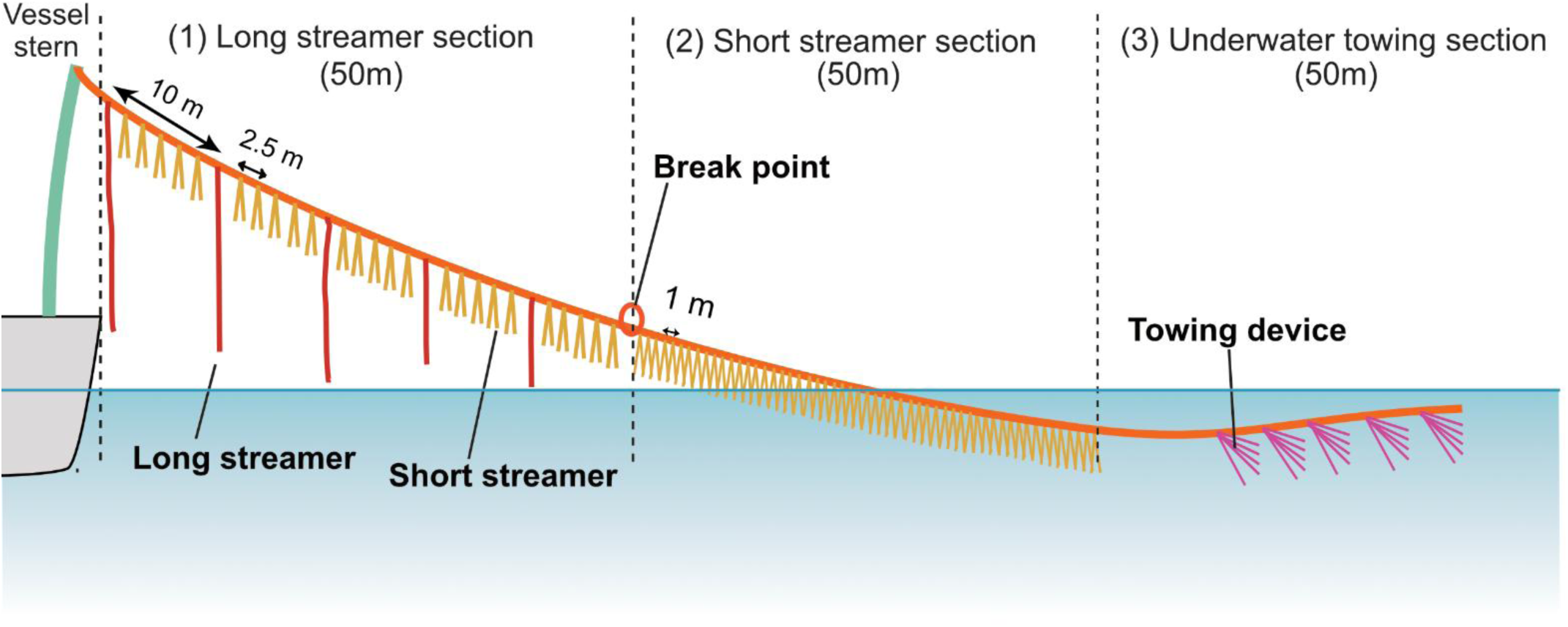
Schematic illustration of the tori-line configuration

**Table 1.** Specification of the hybrid tori-line used in the experiment, including the configuration and dimensions of the long-streamer, short-streamer, and underwater segments.

|  | Specification |
|---|---|
| Main line length | 150m |
| Main line material and color | Polyester multifilament strand, orange |
| Hybrid streamer segment |  |
| Length of the segment | 50m |
| Number of streamers | 50 (long streamer: 5, short streamer: 45) |
| Streamer materials and colors | Long streamer: two folded Nylon code, yellow<br>Short streamer: two folded PP band, yellow |
| Streamer length | Long streamer: 4 m×3 m, 3 m×2<br>Short streamer: 1 m×5 |
| Streamer interval | 2.5m, long and short streamers alternatively deployed |
| Short streamer segment |  |
| Number of streamers | 50 |
| Streamer material and color | Two-folded PP band, yellow |
| Streamer length | 1 m |
| Streamer interval | 1 m |
| Underwater segment |  |
| Towing device | Combined five PP bands deployed into the main line at 5m intervals |

Individual branch lines consisted of a 2 m wire leader attached to a 4.0 sun tuna hook, followed by 8 m of nylon monofilament and 5 m of polyester line (7.2 mm diameter), giving a total branch line length of 15 m (Fig. 1). Weighted branch lines used a double-weight configuration developed and confirmed its effectiveness (Yamazaki’s double weight system; Melvin et al. 2013, 2014, Sato et al. 2013). A 61 g weighted section, consisting of two asymmetric weights (hook side: 26.25 g; branch-line side: 15 g) attached at each end of a 1.5 m nylon-coated lead line (19.75g), was inserted into the monofilament section above the wire leader (Fig. 2).

**Figure 2.**
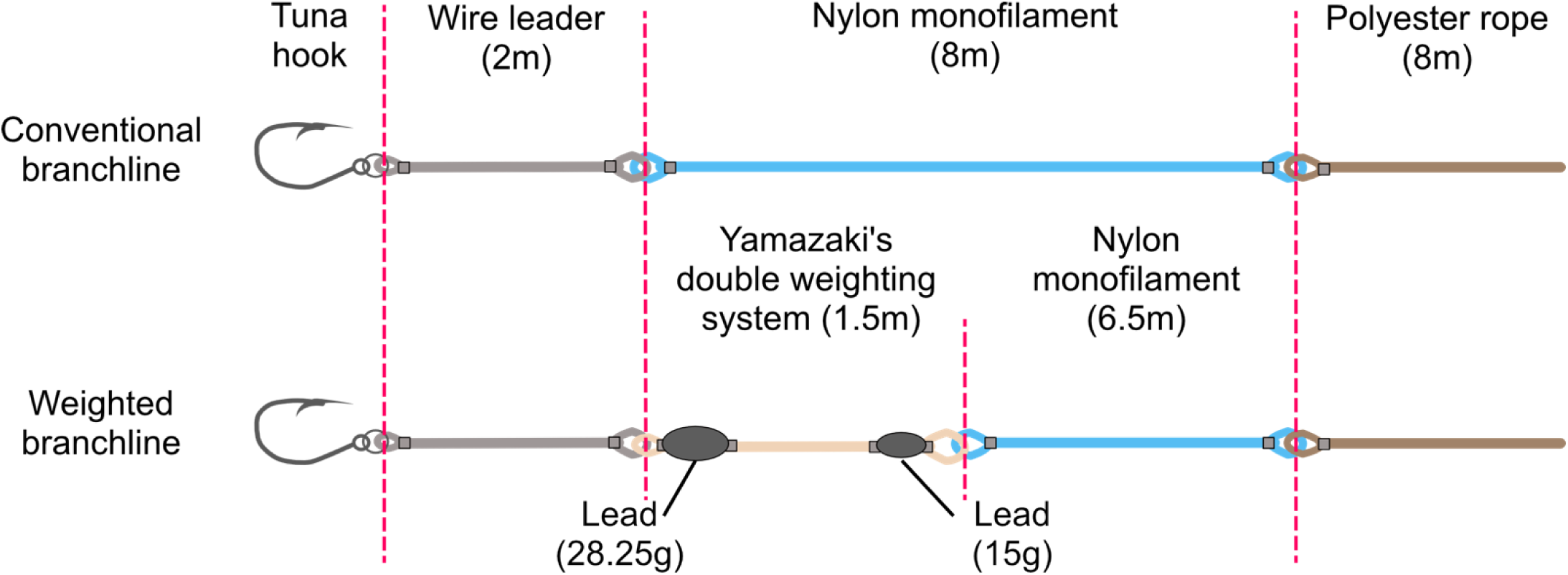
Schematic illustration of unweighted and weighted branch line configurations. The weighted branch line includes a 61 g double-weight section inserted into the monofilament section above the wire leader.

Each operation was divided into six experimental blocks of 160 hooks (40 baskets). Six treatment combinations, consisting of double, single, or no tori-line crossed with weighted or unweighted branch lines, were deployed in each operation (Table 2). After each tori-line treatment change, a 200 m blank section of mainline was inserted to reduce carry-over effects from the previous treatment. Weighted and unweighted branch lines were alternated every 40 hooks, and each tori-line treatment was alternated every 320 hooks, yielding 160 hooks per tori-line × weighting treatment.

**Table 2.** Number of hooks allocated to each experimental block for the six treatment combinations of tori-line deployment (none, single, double) and branch-line weighting (unweighted, weighted).

| Tori line | Weighted branchline |  | Total |
|---|---|---|---|
|  | Unweighted | Weighted |  |
| Double | 160 hooks | 160 hooks | 320 hooks |
| Single | 160 hooks | 160 hooks | 320 hooks |
| None | 160 hooks | 160 hooks | 320 hooks |
| Total | 480 hooks | 480 hooks | 960 hooks |

Seabird interactions were observed during line setting and hauling. Six observation sessions of 20 min each were conducted during each set. During the first 5 min of each session, the number of seabirds attending the operation was recorded and birds were identified to species whenever possible. During the remaining 15 min, seabird attacks on baited hooks were recorded by species and by distance astern in seven classes: 0–25, 26–50, 51–75, 76–100, 101–125, 126–150, and 151–200 m from the stern. Attacks were defined as the initial attempt by an individual bird to retrieve bait from a sinking hook. Secondary attacks, in which one or more birds seize bait already retrieved by another bird, have been identified as an important driver of albatross bycatch in southern hemisphere fisheries, where diving Procellaria petrels bring baited hooks to the surface and surface-foraging albatrosses subsequently take the bait (Melvin et al., 2013, 2014). This mechanism creates a marked mismatch between the species making the primary attack and the species ultimately bycaught. In our study area, diving Procellaria petrels were absent and both focal species were themselves surface foragers, so this source of confounding was unlikely to be substantial. We therefore focused on primary attacks as the relevant behavioural indicator of bycatch risk in this system; detecting secondary attacks also requires a higher level of observer expertise than detecting primary attacks. During observations, attacks were distinguished based on whether they occurred on weighted or unweighted branch lines in the no-tori-line blocks of 29 operations. Wind speed, swell height, and cloud cover were recorded at the start of each observation session. During hauling, the number of bycaught birds was recorded in the experimental blocks where they occurred.

To quantify sinking speed, TDRs (G5, Cefas Technology Limited; DST centi-EX, Star-Oddi Limited) were attached to the junction between the wire leader and the branch line in each operation. The TDRs recorded depth every 1 s at resolutions of 0.01 m (G5) or 0.1 m (DST centi-EX).

Seventeen seabird species were recorded attending line-setting operations during the study period (Table S1). However, bycatch was observed only for Laysan albatross (*Phoebastria immutabilis*) and Black-footed albatross (*P. nigripes*). Species-specific analyses of attack behaviour and bycatch were therefore restricted to these two species. This restriction was based on the occurrence of observed bycatch events, because estimation of the relationship between attack behaviour and bycatch was not possible for species with no recorded captures. Both species are also among the seabirds most frequently incidentally captured in North Pacific pelagic longline fisheries, making them directly relevant focal species for evaluating mitigation performance in the region (Yokota et al. 2007; Sato et al. 2012).

### 2) Statistical analysis

To estimate the effects of tori-lines and weighted branch lines on seabird bycatch mitigation, a two-stage integrated Bayesian model was constructed. Stage 1 described the spatial distribution of seabird attacks relative to the tori-line aerial extent (The area from the stern to where the line attached to the pole enters the water) during line setting. Stage 2 linked the attack rates estimated in Stage 1 with observed bycatch counts in each experimental block. Estimation uncertainty was propagated between stages by passing the Stage 1 posterior predictive distribution into Stage 2. Table 3 provides explanations of the symbols used for the parameter data in Stages 1 and 2.

**Table 3.**
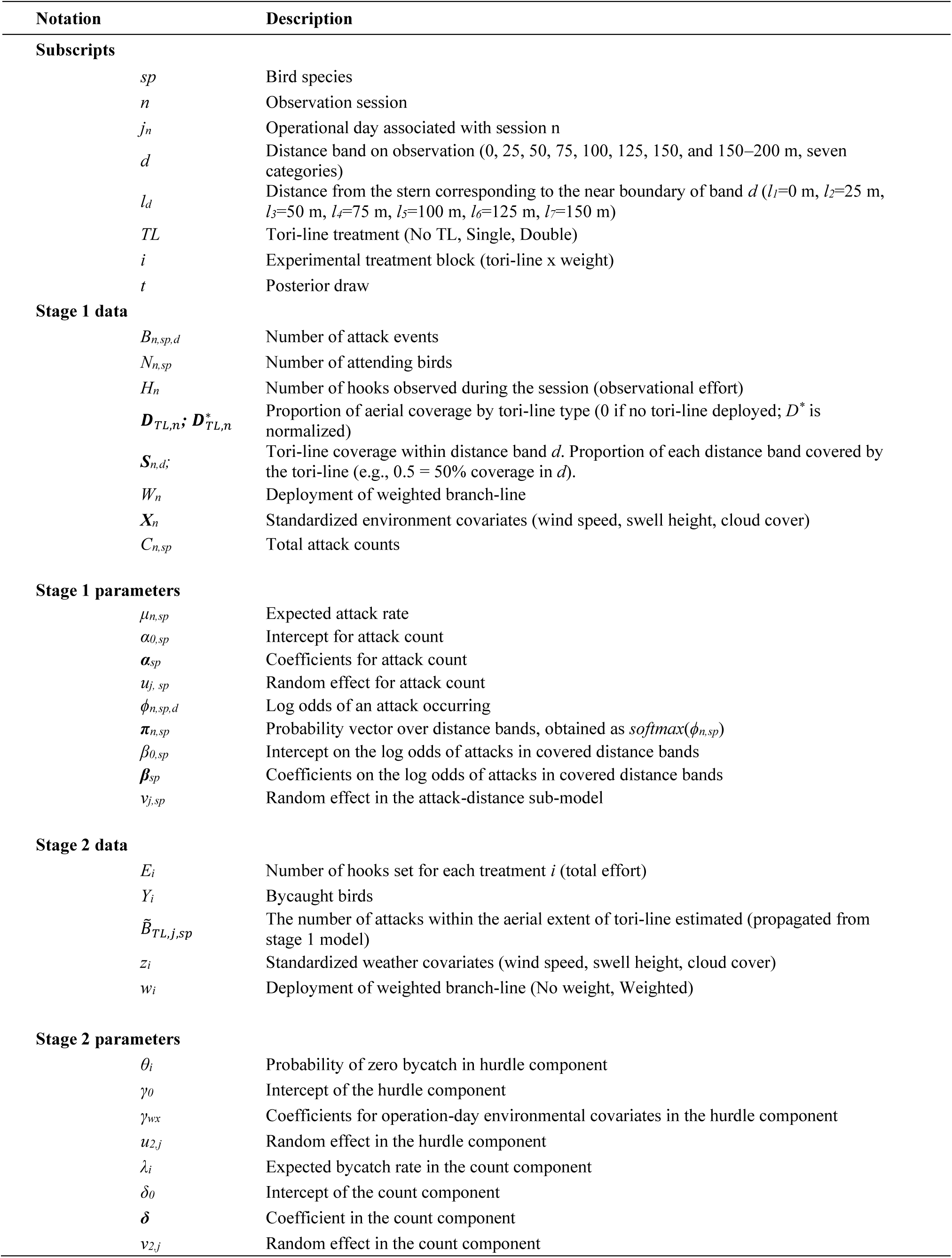
Glossary of notation used in the two-stage integrated Bayesian model.

| Notation | Description |
|---|---|
| **Subscripts** |  |
| *sp* | Bird species |
| *n* | Observation session |
| *j<sub>n</sub>* | Operational day associated with session n |
| *d* | Distance band on observation (0, 25, 50, 75, 100, 125, 150, and 150–200 m, seven categories) |
| *l<sub>d</sub>* | Distance from the stern corresponding to the near boundary of band *d* (*l<sub>1</sub>*=0 m, *l<sub>2</sub>*=25 m, *l<sub>3</sub>*=50 m, *l<sub>4</sub>*=75 m, *l<sub>5</sub>*=100 m, *l<sub>6</sub>*=125 m, *l<sub>7</sub>*=150 m) |
| *TL* | Tori-line treatment (No TL, Single, Double) |
| *i* | Experimental treatment block (tori-line x weight) |
| *t* | Posterior draw |
| **Stage 1 data** |  |
| *B<sub>n,sp,d</sub>* | Number of attack events |
| *N<sub>n,sp</sub>* | Number of attending birds |
| *H<sub>n</sub>* | Number of hooks observed during the session (observational effort) |
| ***D***<sub>*TL,n*</sub>***;*** ***D***<sup>\*</sup><sub>*TL,n*</sub> | Proportion of aerial coverage by tori-line type (0 if no tori-line deployed; *D*<sup>\*</sup> is normalized) |
| ***S***<sub>*n,d;*</sub> | Tori-line coverage within distance band *d*. Proportion of each distance band covered by the tori-line (e.g., 0.5 = 50% coverage in *d*). |
| *W<sub>n</sub>* | Deployment of weighted branch-line |
| ***X***<sub>*n*</sub> | Standardized environment covariates (wind speed, swell height, cloud cover) |
| *C<sub>n,sp</sub>* | Total attack counts |
| **Stage 1 parameters** |  |
| *μ<sub>n,sp</sub>* | Expected attack rate |
| *α<sub>0,sp</sub>* | Intercept for attack count |
| ***α***<sub>*sp*</sub> | Coefficients for attack count |
| *u<sub>j, sp</sub>* | Random effect for attack count |
| *ϕ<sub>n,sp,d</sub>* | Log odds of an attack occurring |
| ***π***<sub>*n,sp*</sub> | Probability vector over distance bands, obtained as *softmax*(*ϕ<sub>n,sp</sub>*) |
| *β<sub>0,sp</sub>* | Intercept on the log odds of attacks in covered distance bands |
| ***β***<sub>*sp*</sub> | Coefficients on the log odds of attacks in covered distance bands |
| *v<sub>j,sp</sub>* | Random effect in the attack-distance sub-model |
| **Stage 2 data** |  |
| *E<sub>i</sub>* | Number of hooks set for each treatment *i* (total effort) |
| *Y<sub>i</sub>* | Bycaught birds |
| *B̃<sub>TL,j,sp</sub>* | The number of attacks within the aerial extent of tori-line estimated (propagated from stage 1 model) |
| *z<sub>i</sub>* | Standardized weather covariates (wind speed, swell height, cloud cover) |
| *w<sub>i</sub>* | Deployment of weighted branch-line (No weight, Weighted) |
| **Stage 2 parameters** |  |
| *θ<sub>i</sub>* | Probability of zero bycatch in hurdle component |
| *γ<sub>0</sub>* | Intercept of the hurdle component |
| *γ<sub>wx</sub>* | Coefficients for operation-day environmental covariates in the hurdle component |
| *u<sub>2,j</sub>* | Random effect in the hurdle component |
| *λ<sub>i</sub>* | Expected bycatch rate in the count component |
| *δ<sub>0</sub>* | Intercept of the count component |
| ***δ*** | Coefficient in the count component |
| *v<sub>2,j</sub>* | Random effect in the count component |

#### Stage 1

The observational unit in Stage 1 was the behavioral observation session during line setting. Multiple sessions were conducted on each operation day, and predictor values varied at the session level. Total attacks were defined as

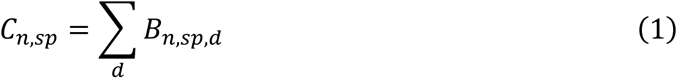

Stage 1 took the form of a finite-mixture model with two sub-models fitted simultaneously.

The first sub-model described total attack count using a Poisson regression:

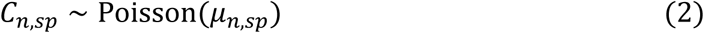

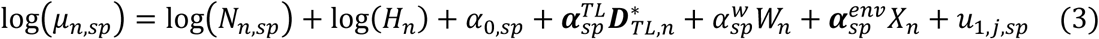

The offset term log(*N_n_*_,*sp*_) + log(*H_n_*) normalized the expected count by the number of attending birds and the number of observed gear units, and *u_j_*_(*n*),*sp*_was a species-specific random effect for operation day.

The second sub-model described the spatial distribution of attacks using a multinomial regression:

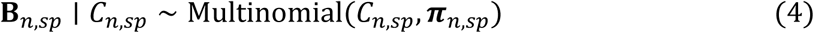

For *d* = 1, … ,6;

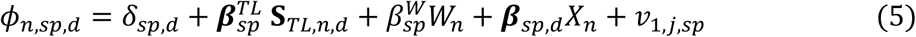

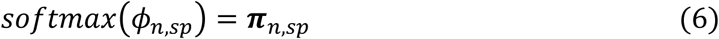

with the reference category *d* = 7 (beyond 150 m) fixed as *φ_n_*_,*sp*,7_ = 0, and *v*_1,*j*(*n*),*sp*_ was a random effect for operation day. Sessions with zero attacks contributed only to the Poisson likelihood. To improve computational efficiency, random effects in both sub-models were parameterized in non-centered form.

#### Stage 2

A quantity was extracted from the Stage 1 posterior predictive distribution for use in Stage 2. First, for each posterior sample, the number of attacks within the effective tori-line range was aggregated across sessions for each operation day:

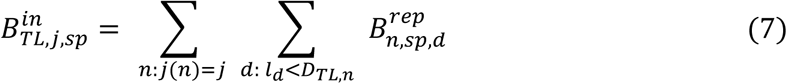

where 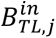 represents the total number of attacks occurring within the effective aerial coverage of the tori-line on operation day *j*, aggregated across sessions and computed from posterior predictive replicate 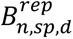 of Stage 1. Sessions without tori-lines were excluded. The corresponding log-scale variable used in Stage 2 was

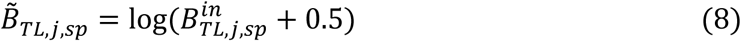

where 0.5 was added to avoid log(0).

Stage 2 modeled bycatch counts recorded for each experimental block using a Poisson hurdle model, which separated the occurrence of any bycatch from the number of birds caught when bycatch occurred. Mitigation measures were included only in the count component under the mechanistic assumption that environmental and operational conditions mainly determine whether bycatch occurs at all, whereas mitigation devices influence capture probability after bird–gear contact.

Because Black-footed albatross bycatch was zero in all tori-line treatment blocks, this species was analyzed in a separate sub-model restricted to the no-tori-line treatments, in which only the weight effect was estimated. Laysan albatross was analyzed in the main model including all treatment combinations.

For both models, the hurdle component was specified as

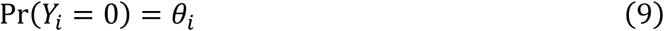

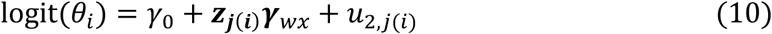

where *θ_i_* is the probability of no bycatch in block *i*, and *u*_2,*j*(*i*)_ is a random effect for the hurdle component.

The count component for the Laysan albatross model was

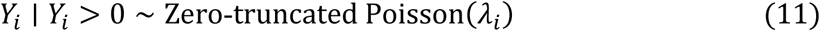

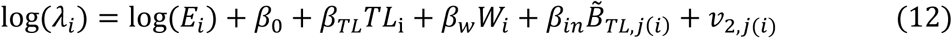

The Black-footed albatross model, fitted only to the no-tori-line treatment, was

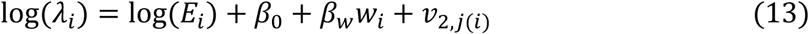

The offset term log(*E_i_*) corrected for hook number. Random intercepts for the hurdle and count components were fitted independently using non-centered parameterization.

All models were implemented in Stan (Stan Development Team 2024) and fitted with Hamiltonian Monte Carlo through the *cmdstanr* package in R (Gabry and Češnovar 2022). Four chains of 5,000 iterations were run, with the first 1,000 iterations in each chain discarded as warm-up, leaving 16,000 posterior samples per parameter. Convergence was assessed using rank-normalized split-*R*^^^ and effective sample size (Vehtari et al. 2021).

#### Evaluation of estimated effects

The effects of mitigation treatments were evaluated from posterior predictions after marginalizing over the remaining parameters. In Stage 1, predicted values were obtained for total attack count and attack-distance distribution. In Stage 2, predicted bycatch counts were calculated for each mitigation treatment, and the estimated coefficients for mitigation measures and within-range attack counts were also examined. These predicted values and log rate ratios were calculated from the expected values obtained by combining the hurdle component and the count component.

The principal effect-size measure was summarized as posterior predictive log rate ratios relative to the no-mitigation control, an effect-size metric equivalent to the log response ratio widely used in ecological research (Hedges et al., 1999; Lajeunesse, 2015).

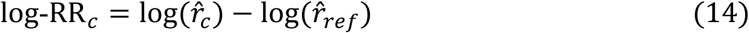

Posterior inference was based on a Region of Practical Equivalence (ROPE) framework (Kruschke 2018; Kruschke and Liddell 2018). For log-scale parameters representing total attack count and bycatch rate, the ROPE width was set at ±log(1.1), corresponding to a ±10% change in rate. For logit-scale parameters describing distance distribution, the ROPE width was set at ±0.2. The 95% highest density interval was classified as a substantial decrease, practical equivalence, substantial increase, or inconclusive depending on its relationship to the ROPE boundaries. Effects were classified by comparing the 95% HDI of each posterior distribution with the corresponding ROPE (Kruschke, 2018; Kruschke and Liddell, 2018). An effect was judged as practical equivalence when the 95% HDI lay entirely within the ROPE, and as a substantive decrease or substantive increase when the 95% HDI lay entirely outside the ROPE on the negative or positive side, respectively. When the 95% HDI partially overlapped the ROPE, the effect was classified as inconclusive, because the data did not allow the two possibilities to be distinguished with the specified precision. In the figures, this classification corresponds to whether the horizontal bar (95% HDI) is fully inside the shaded ROPE band, fully outside it on one side, or straddles its boundary.

## 3. Results

Weighted branch lines reached depths of 2 and 5 m at mean sink rates of 0.18 and 0.21 m s⁻¹, respectively, compared with 0.14 and 0.17 m s⁻¹ for unweighted branch lines. Thus, weighted branch lines sank significantly faster than unweighted branch lines at both depths (Gamma GLM, N = 591, p < 0.001).

All integrated models converged successfully. The *R*^^^ values for all parameters were below 1.01, and all effective sample sizes exceeded 400.

Both tori-lines and weighted branch lines contributed to reducing albatross overall attack counts. However, there were species differences by each mitigation gear. For Laysan albatross, there was a tendency for the reduction associated with double tori-lines to be smaller than that associated with a single tori-line while it was same level between both for Black-footed albatross (Fig. 3).

**Figure 3.**
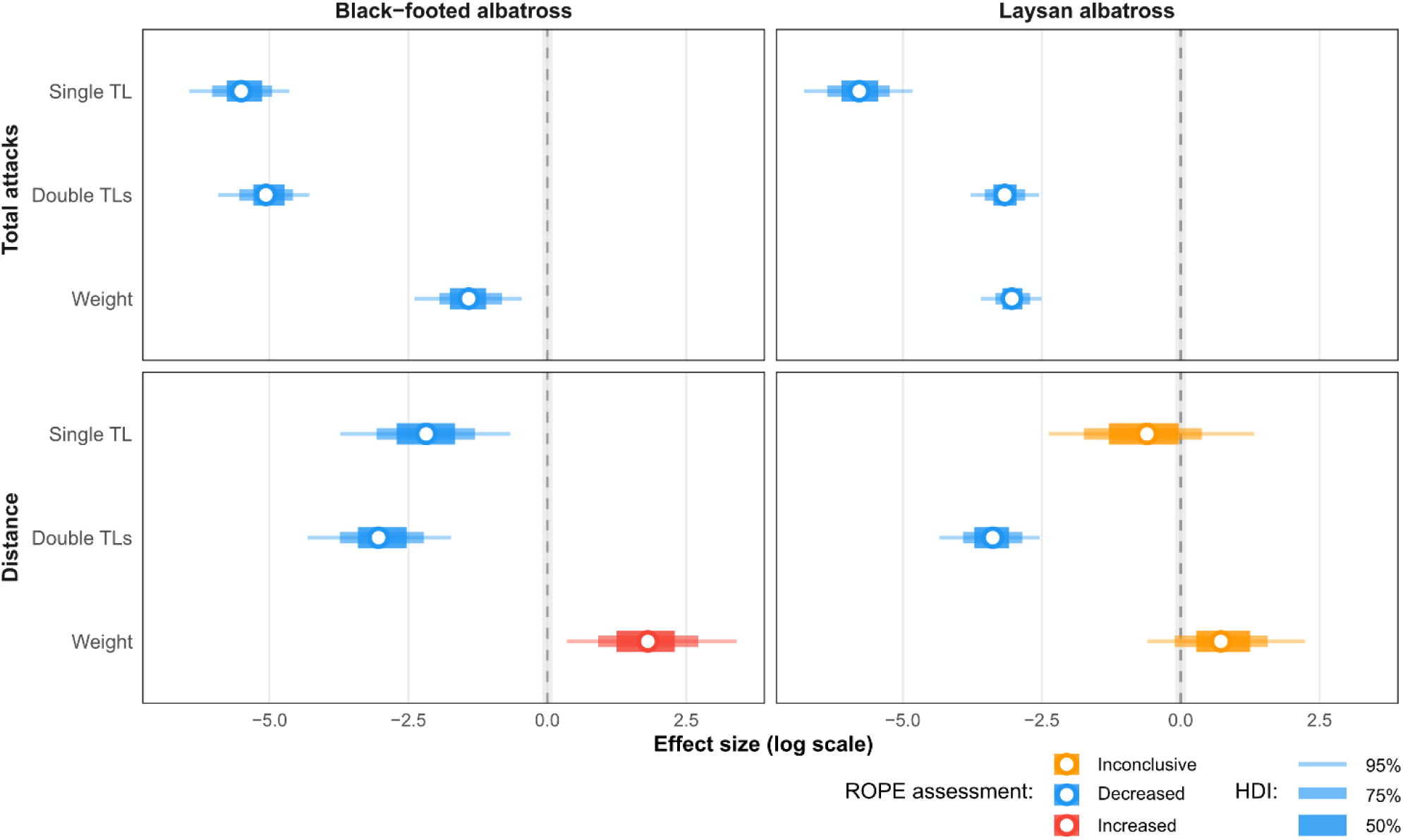
Posterior estimates of the fixed effects of mitigation treatments in Stage 1 of the integrated Bayesian model for total attack count and attack-distance distribution, shown separately for Black-footed albatross and Laysan albatross. Points indicate posterior medians and horizontal bars indicate 95% highest density intervals. The shaded region indicates the ROPE.

The effect of mitigation gear on the attack-distance model component showed contrasting responses between tori-lines and weighted branch lines, with species-specific differences also evident. For Black-footed albatross, weighted branch lines were associated with attacks occurring closer to the vessel, whereas for Laysan albatross, only the double tori-line clearly shifted attacks farther from the vessel and the effects of a single tori-line and of weighted branchline on attack distance were inconclusive.

Posterior predictive values of total attack counts showed that all mitigation treatments reduced total attacks and that combining devices produced additive reductions in attack count (Fig. 4). For Black-footed albatross, the combination of tori-line and weighted branch line gave the largest reduction in attacks while weighted branch line alone had the weakest effect. For Laysan albatross, the largest reduction was obtained with the combination of a single tori-line and weighted branchline while double tori-lines produced the smallest attack reduction.

**Figure 4.**
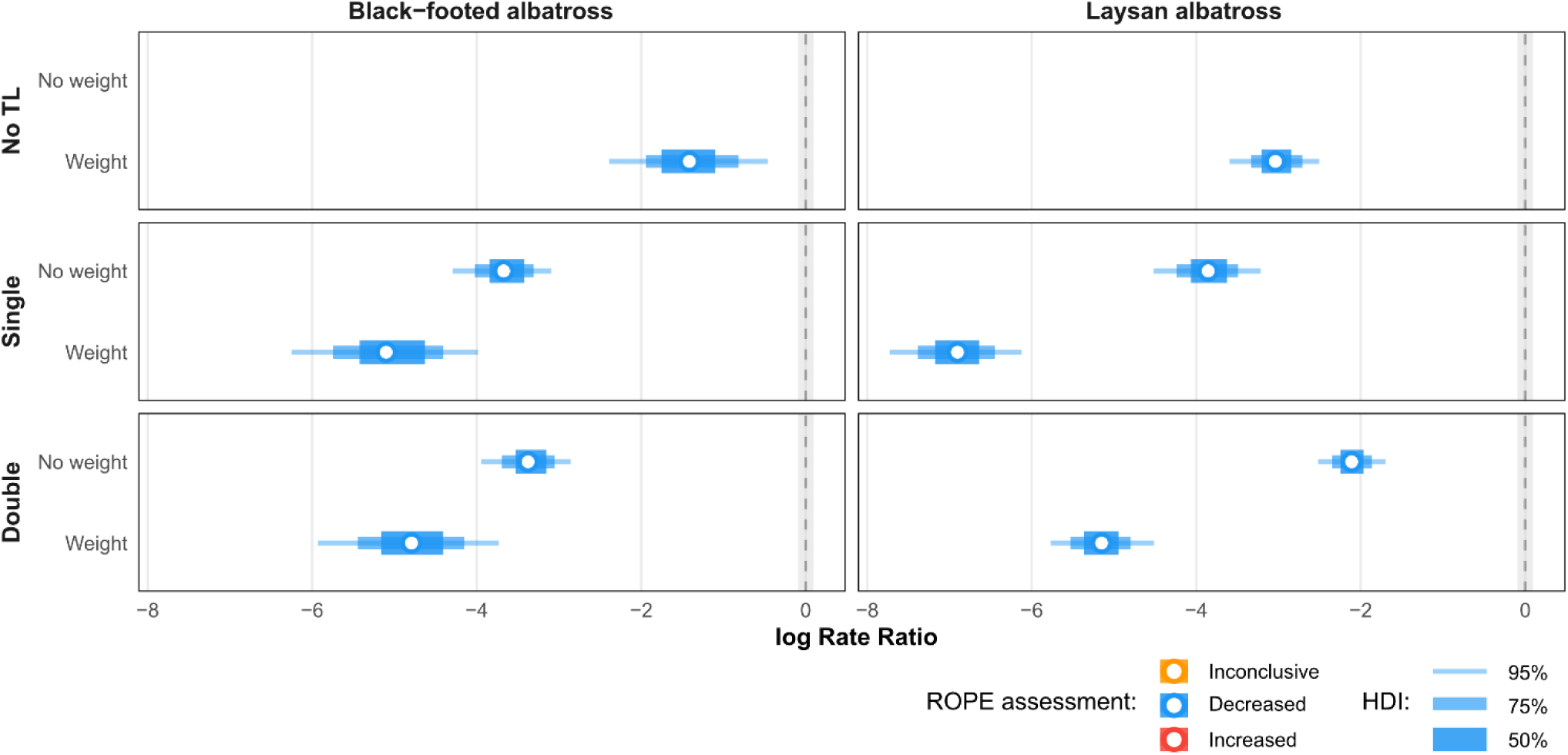
Posterior predictive log rate ratios of total attack counts for each mitigation treatment combination relative to the control condition without mitigation devices, derived from Stage 1 of the integrated Bayesian model. Points indicate posterior medians and horizontal bars indicate 95% highest density intervals. The shaded region indicates the ROPE. No value is shown for the unweighted, no-tori-line combination because it is the reference condition against which all other treatments were compared.

Posterior predictive values of attack count within and beyond the effective range of the tori-line showed that double tori-lines clearly reduced attacks within the range while increasing attacks beyond it, whereas weighted branch lines contributed little to attack counts on either side of the range (Fig. 5). Species differences were nonetheless evident in the single tori-line treatment. For Black-footed albatross, single tori-line increased attacks beyond the effective range, whereas changes within the range were unclear. For Laysan albatross, attack position changed little with a single tori-line.

**Figure 5.**
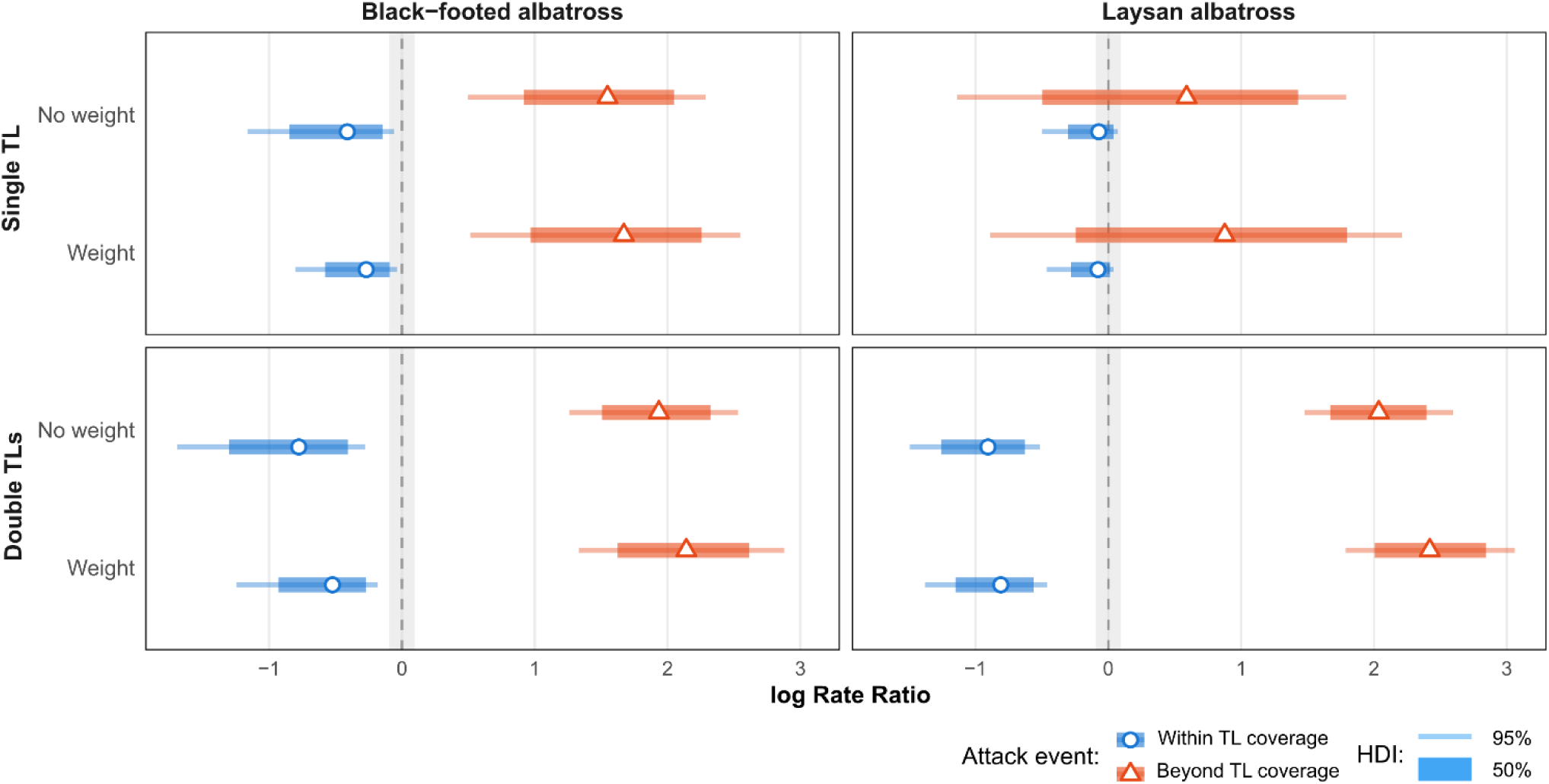
Posterior predictive log rate ratios of attack counts occurring within and beyond the effective range of the tori-line for each mitigation treatment combination relative to the control condition, derived from Stage 1 of the integrated Bayesian model. Points indicate posterior medians and horizontal bars indicate 95% highest density intervals. The shaded region indicates the ROPE.

In the Stage 2 model, all tori-line treatments contributed to reducing Laysan albatross bycatch, although the contribution of weighting was smaller for both Laysan and Black-footed albatross (Fig. 6). The within-range attack frequency estimated in Stage 1 was clearly associated with an increase in Laysan albatross bycatch. For Black-footed albatross, the estimated effect of weighted branch line in the no-tori-line submodel was negative but not clearly distinguishable from zero.

**Figure 6.**
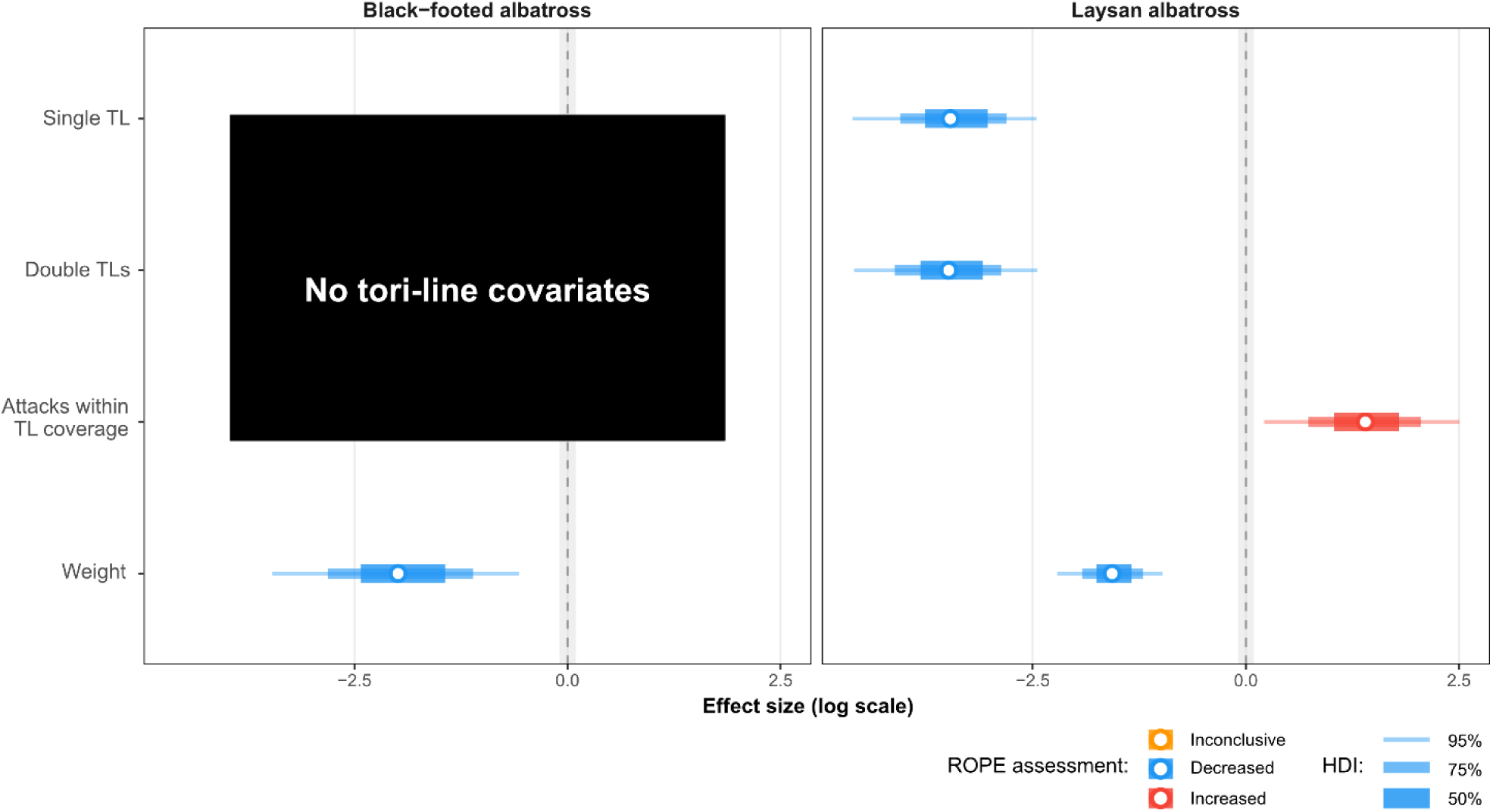
Posterior estimates of fixed effects in Stage 2 of the integrated Bayesian model for bycatch count, including mitigation treatments and the contribution of attacks within the effective tori-line range. Points indicate posterior medians and horizontal bars indicate 95% highest density intervals. The shaded region indicates the ROPE.

Posterior predicted value of bycatch counts showed substantive reductions for Laysan albatross under all mitigation treatments, but differences among treatment combinations were small and generally inconclusive (Fig. 7). Additive effect by combination of devices that was found in attack count were not observed. For Black-footed albatross, observed bycatch was zero in all tori-line treatment blocks. Accordingly, the model-based predictions for this species were restricted to the no-tori-line treatments, and the effect of weighted branchline remained inconclusive.

**Figure 7.**
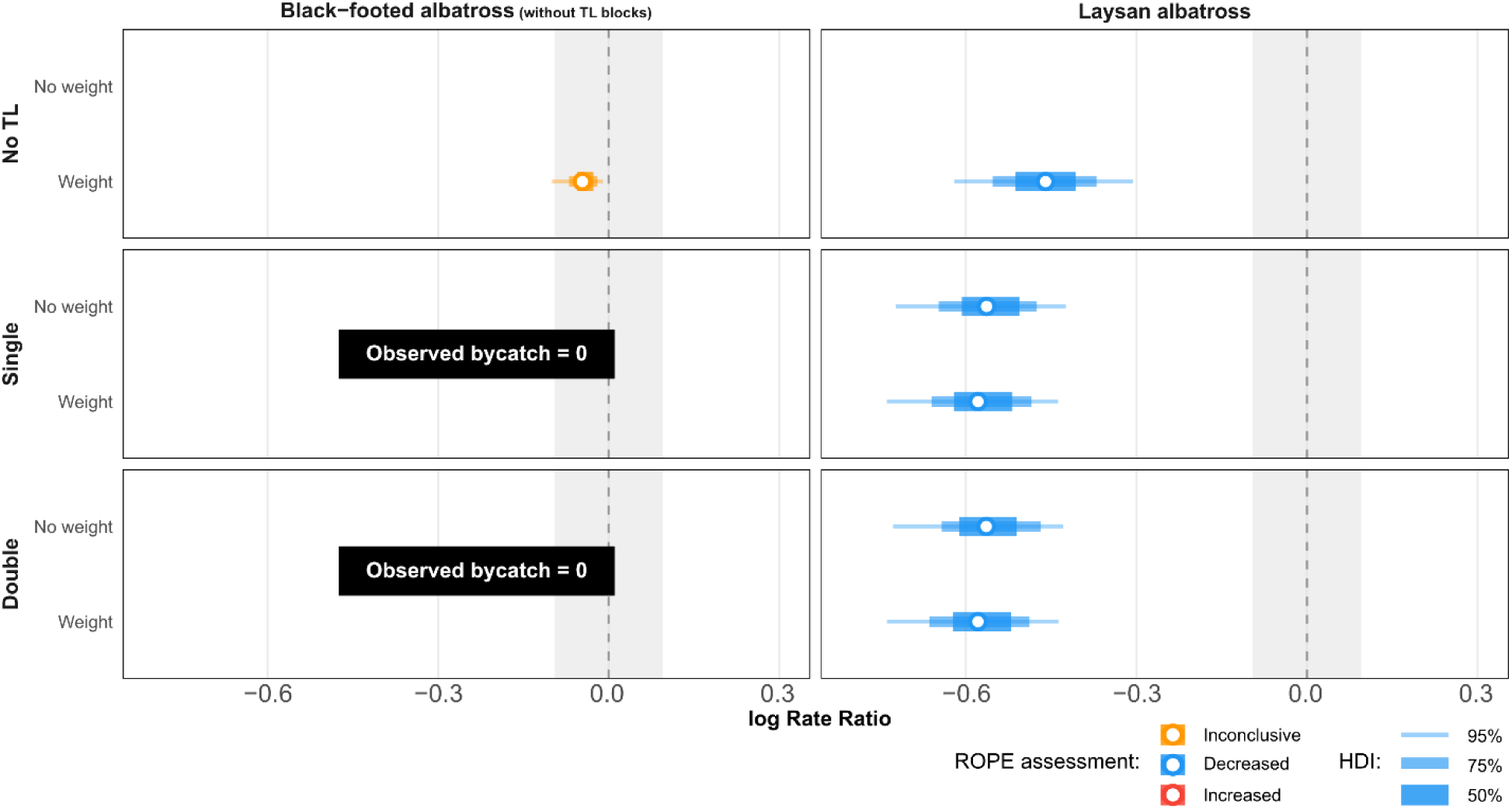
Posterior predictive log rate ratios of bycatch counts for each mitigation treatment combination relative to the control condition, derived from Stage 2 of the integrated Bayesian model. Points indicate posterior medians and horizontal bars indicate 95% highest density intervals. The shaded region indicates the ROPE. For Black-footed albatross, predictions are restricted to the no-tori-line sub-model because observed bycatch was zero in all tori-line treatment blocks.

## 4. Discussion

This study used multiple metrics derived from two-stage models to demonstrate how tori-lines and weighted branch lines reduce bycatch: tori-line deployment reduced attacks within the aerial coverage zone, where bait remains near the surface, and this reduction was associated with lower bycatch. The number of tori-lines deployed also altered the pattern of attack-distribution. Deploying two tori-lines produced an additional “stand-off” effect—a shift in attack location away from the vessel and beyond the aerial coverage zone—whereas weighted branch lines reduced attack frequency without pushing attacks away from the coverage zone and, for one species, increased the probability that attacks occurred within the zone. Because this experiment was conducted in an area with few diving seabirds, the species differences observed here cannot be attributed to differences in diving capability. Nonetheless, clear species differences were evident: the stand-off effect of a single tori-line was weak for Laysan albatross, while tori-line deployment produced a strong reduction in bycatch for Black-footed albatross. These results provide information relevant to tori-line specifications and to the operation of mitigation measures by regional fisheries management organizations and local fisheries.

The present analysis also made it possible to separate two components of deterrence during line setting: a reduction in the absolute number of attacks, and a stand-off effect that displaces attack location beyond the aerial coverage zone of the tori-line. This dual perspective allows a more detailed evaluation of both the process generating attack behavior and its suppression. Most of previous studies have discussed mitigation performance in terms of reduced bait-taking attacks (Sato et al. 2012; Melvin et al 2013; Gilman et al. 2021), but the present results suggest that overall attack suppression and the stand-off effect are distinct processes.

Tori-lines consistently reduced total attacks and, in several cases, produced a stand-off effect. A notable finding was that using double tori-lines, unlike a single line, produced a clear stand-off effect, pushing attacks farther from the vessel. For example, Sato et al. (2013) and Cortés and González-Solís (2018) reported a significant reduction in attacks within the tori-line’s aerial coverage area but did not detect a stand-off effect. Deploying two tori-lines likely reduced the opportunity for birds to enter the aerial coverage zone in the first place. This may partly indicate that birds redirected their attacks to areas beyond the end of tori-line aerial coverage, but Zhou and Brothers (2022) showed that bycatch risk is unlikely to be equal across attack attempts and may vary with operational conditions and seabird assemblage. Operating two tori-line units may make it difficult for seabirds to pinpoint the location of the bait, which could make it harder for them to target and attack the bait even beyond the end of aerial coverage.

By contrast, weighted branch lines reduced total attack frequency without producing a stand-off effect, a pattern consistent with the expectation that faster sink rates reduce bait availability rather than altering where birds attempt to attack. This overall reduction in attack frequency is consistent with previous research on weighted branch lines (Robertson et al. 2013; Melvin et al. 2014). However, because weighted branch lines alone do not prevent birds from attacking hooks before bait has sunk beyond their reach, this measure alone is insufficient to eliminate high-risk attacks near the vessel. Combining weighted branch lines with tori-lines or other measures that keep birds away from the vessel is therefore important for effective bycatch mitigation.

By propagating the Stage 1 posterior distribution of within-range attack counts into the bycatch model (Hooten and Hefley 2019), the effect of mitigation measures on bycatch could be evaluated while directly accounting for attack frequency. For Laysan albatross, attack frequency within the tori-line coverage zone contributed directly to increased bycatch, supporting previous suggestions (Sato et al. 2013) and, to our knowledge, providing the first explicit demonstration of a direct relationship between attack behavior and bycatch events. This result partially validates the use of attack counts under tori-line coverage as a proxy for bycatch risk, an assumption that has previously been applied without direct confirmation and that underlies much of the mitigation-gear evaluation literature. Because attack counts can be recorded far more frequently than actual bycatch events, and without requiring birds to be captured or killed, a validated behavioral proxy would allow mitigation performance to be screened under a much wider range of operational conditions than lethal-outcome experiments alone permit.

Bycatch was reduced by any single mitigation device except weighted branch lines for Black-footed albatross, with tori-line effectiveness similar regardless of line number and little additional benefit from combining devices with weighting. The similarity across tori-line configurations, despite the stand-off effect being detected only for double lines, suggests a ceiling effect: once a tori-line suppresses most attacks within its coverage area, further displacing attacks beyond that area adds little additional protection, because the seabirds it would displace are already deterred from attacking. The limited additive benefit of weighting is consistent with this interpretation, as weighted branch lines act on bait accessibility through sink rate rather than on bird behavior, addressing a risk that tori lines have already largely removed. This also helps explain the species-specific shortfall for Black-footed albatross: the weighted branch line configuration used here was validated in southern hemisphere waters with abundant diving seabirds (Melvin et al. 2013; Düssler et al. 2026), whereas North Pacific seabirds, including the species examined here, have comparatively limited diving capability (Matsumoto et al. 2012; Kazama et al. 2019) and tend to attack near the surface (Sato et al. 2012); a device designed to move bait beyond diving range offers correspondingly less protection against species that rarely dive to begin with.

The estimated effects of mitigation devices differed between Black-footed and Laysan albatrosses. Both species showed consistent reductions in total attack rate across treatments, but attack-distance and bycatch responses differed. The differences may reflect species-specific foraging behavior associated with body size and flight style (Conners et al. 2015). This result supports previous suggestions that regional differences in seabird assemblages influence both bycatch mechanisms and the effectiveness of mitigation measures (Sato et al. 2012; Zhou and Brothers 2022). Because the performance of mitigation devices depends on the foraging behavior of the species exposed to the fishery, measures validated in other regions should not be assumed to perform equally well elsewhere.

Our results indicated that tori-lines were the most effective mitigation device in the western North Pacific, the region examined in this study. Weighted branch lines showed some evidence of effectiveness for Laysan albatross, but not clearly for Black-footed albatross. Predicted effect sizes for weighted treatments differed little from those for tori-line-only treatments, suggesting that weighted branch lines were less effective overall than tori-lines. The weight configuration used here was the same as that validated in southern hemisphere studies, making the regional difference in performance noteworthy. Recognizing that the optimal mitigation device differs by region is important: seabird assemblages with different foraging modes, and fisheries with different operational styles, occur in different regions, and the combination of measures that balance feasibility in operational circumstance and mitigation effectiveness should therefore differ accordingly. The quantitative evaluation framework developed here can support decision-making in selecting mitigation measures suited to each region.

### 1) Implications for Evaluating and Implementing Seabird Bycatch Mitigation Measures

Future studies should report the operational characteristics that determine realised tori-line performance, including the number and side of deployed lines, effective aerial extent under prevailing conditions, and loss of coverage caused by entanglement or breakage. Such information is needed to allow meaningful comparisons among studies and fisheries. Tori-line configurations should be evaluated in relation to the local seabird assemblage and the sinking performance of baited hooks. In particular, differences in species’ tendency to forage near vessels and their diving capability should be considered when selecting the extent and configuration of aerial coverage.

## Acknowledgements

The authors thank E.F. Melvin and G. Troy for their assistance with experimental design and research equipment; the crew of the Taikei-Maru No. 2, particularly the skipper Asano, O. for their helpful advice on arranging the survey plan; H. Matsunaga and K. Yokawa, members of the bycatch group at the Shimizu Office; as well as other collaborators, Yashige, R., involved in the research cruise. This research was conducted and funded as a part of the research and assessment program for internationally managed fisheries resources, the Fisheries Agency of Japan. The authors used Perplexity (https://www.perplexity.ai/) as a Generative AI to assist with English writing. The authors reviewed and edited all AI-generated suggestions as needed and accept full responsibility for the content of the publication.

---

**Table S1.**
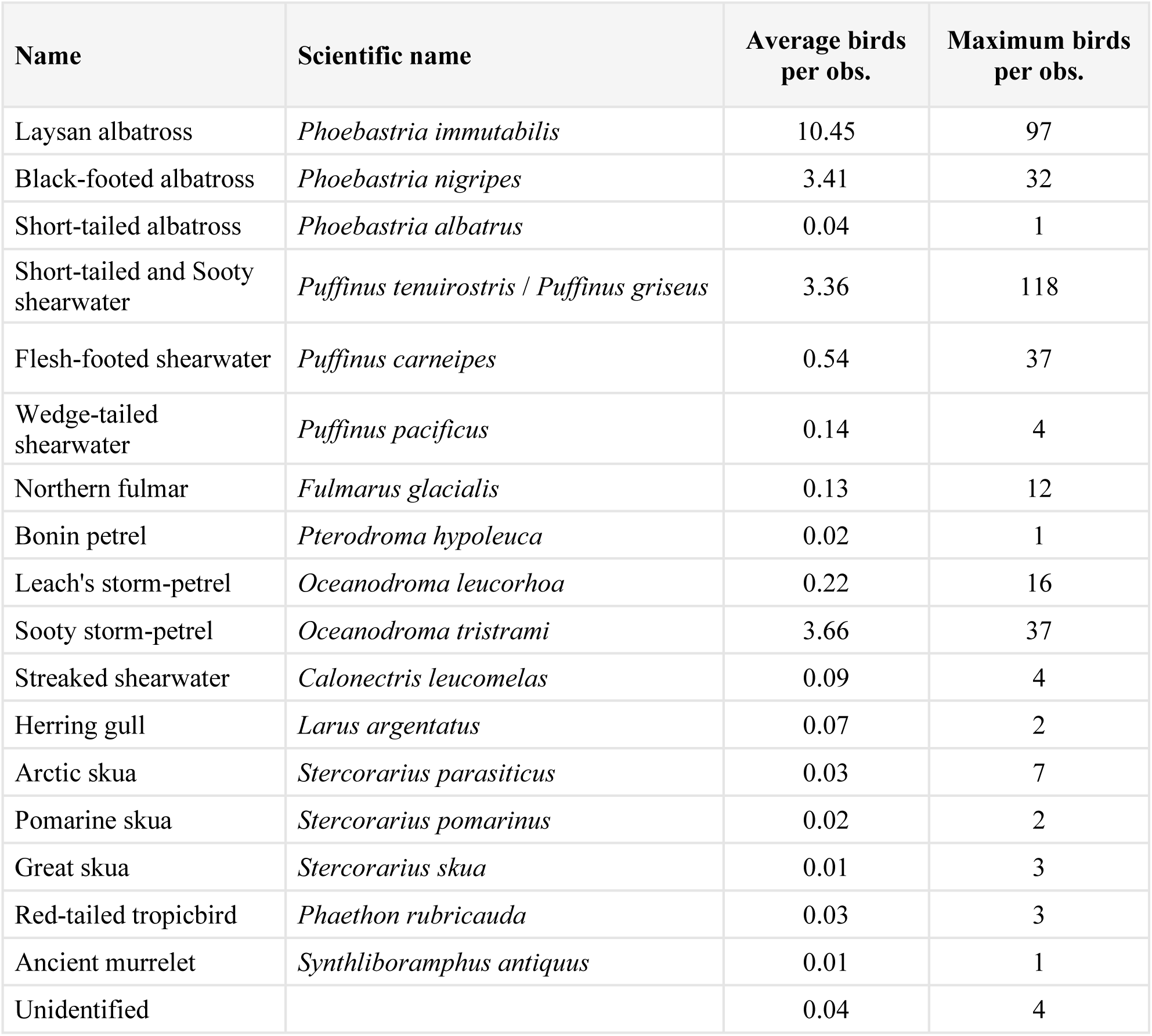
List of seabirds observed during the operational experiment.

